# Caspase-resistant ROCK1 expression prolongs survival of *Eµ-Myc* B cell lymphoma mice

**DOI:** 10.1101/2024.02.26.582074

**Authors:** Katerina Mardilovich, Gregory Naylor, Linda Julian, Narisa Phinichkusolchit, Karen Keeshan, Karen Blyth, Michael F Olson

**Affiliations:** Cancer Research UK Beatson Institute, Garscube Estate, Switchback Road, Glasgow, G61 1BD, UK; Adaptimmune, Abingdon, OX14 4RX, UK; School of Cancer Sciences, University of Glasgow, Glasgow, G16 1QH, UK; Beatson West of Scotland Cancer Centre, NHS Greater Glasgow & Clyde, 1053 Great Western Road, Glasgow, G12 0YN, UK; Cancer Research UK Cambridge Institute, Robinson Way, Cambridge, CB2 0RE, UK; Wolfson Wohl Cancer Research Centre, Paul O’Gorman Leukemia Research Centre, School of Cancer Sciences, College of Medical Veterinary and Life Sciences, University of Glasgow, Glasgow, UK; Toronto Metropolitan University, Department of Chemistry and Biology, 350 Victoria Street, Toronto ON, M5B 2K3, Canada

## Abstract

Apoptosis is characterized by membrane blebbing and apoptotic body formation. Caspase cleavage of ROCK1 generates an active fragment that promotes actin-myosin mediated contraction and membrane blebbing during apoptosis. Expression of caspase-resistant non-cleavable ROCK1 (*Rock1 NC*) prolonged survival of mice that rapidly develop B cell lymphomas due to *Eµ-Myc* transgene expression. *Eµ-Myc; Rock1 NC* mice had significantly fewer bone marrow cells relative to *Eµ-Myc* mice expressing wild-type ROCK1 (*Rock1 WT)*, which was associated with altered cell cycle profiles. Circulating macrophage numbers were lower in *Eµ-Myc; Rock1 NC* mice, but there were higher levels of bone marrow macrophages, consistent with spontaneous cell death in *Eµ-Myc; Rock1 NC* mice bone marrows being more inflammatory. *Rock1 WT* recipient mice transplanted with pre-neoplastic *Eµ-Myc; Rock1 NC* bone marrow cells survived longer than mice transplanted with *Eµ-Myc; Rock1 WT* cells, indicating that the survival benefit was intrinsic to the *Eµ-Myc; Rock1 NC* bone marrow cells. The results suggest that the apoptotic death of *Eµ-Myc; Rock1 NC* cells generates a proliferation-suppressive microenvironment in bone marrows that reduces cell numbers and prolongs B cell lymphoma mouse survival.

## Introduction

The *c-MYC* transcription factor was one of the first proto-oncogenes identified, initially discovered as the cellular version of a viral oncogene associated with avian tumours (Duesberg et al., 1977). Evidence that endogenous *c-MYC* could act as an oncogene came from the discovery of chromosomal translocations between chromosomes 8 and 14 in Burkitt’s lymphoma that placed the *c-MYC* gene adjacent to immunoglobulin heavy chain (IgH) transcriptional enhancer sequences, resulting in elevated *c-MYC* transcription and increased protein levels (Dalla-Favera et al., 1982; Neel et al., 1982; Taub et al., 1982). Experimental evidence for this translocation being a driver of B cell lymphomas was provided by one of the first mouse transgenic cancer models, in which the *c-MYC* gene was coupled to IgH µ enhancer sequences to establish the *Eµ-Myc* mouse line (Adams et al., 1985). These mice rapidly developed lymphomas, with the presence of both immature and mature B lymphocytes indicating that *c-MYC* was oncogenic at various stages of B-cell maturation (Adams et al., 1985). Intriguingly, *c-MYC* activation was found to be a potent inducer of apoptotic cell death in addition to being a driver of cell cycle progression (Evan et al., 1992). In the context of the *Eµ-Myc* mouse model, the high apoptotic rate of MYC expressing cells is associated with the presence of numerous tingible body macrophages containing engulfed TUNEL-positive apoptotic cells within lymphoma tumours (Park et al., 2005). Many studies have examined the interaction of c-MYC with pro- and anti-apoptotic proteins in the induction of B cell lymphomas, including p53, p19^ARF^, BCL-2, BCL-XL, Fas and FasL (reviewed in reference (Morton and Sansom, 2013)). One variable that had not been previously considered is whether the morphological changes associated with apoptotic cell death influences the incidence, characteristics, or severity of *c-MYC*-induced B cell lymphomas.

We previously demonstrated that the caspase-mediated cleavage and consequent hyperactivation of the Rho-associated coiled-coil kinase 1 (ROCK1) is the major driver of the morphological responses observed during apoptotic cell death, including cell contraction, membrane blebbing, apoptotic body formation, and nuclear disintegration (Coleman et al., 2001; Croft et al., 2005). By establishing a genetically modified mouse model in which a single aspartic acid residue in the ROCK1 caspase cleavage site was changed to alanine (D1113A), the absolute dependency of the characteristic apoptotic morphological events on ROCK1 cleavage was demonstrated (Julian et al., 2021). Mouse embryonic fibroblasts (MEFs) expressing non-cleavable ROCK1 (*Rock1 NC*) responded to pro-apoptotic stimuli with typical caspase activation and phosphatidylserine externalization. However, ROCK1 cleavage was blocked, phosphorylation of the regulatory non-muscle myosin light chain was reduced, and cells were significantly impaired in their ability to generate contractile force relative to MEFs expressing wild-type ROCK1 (*Rock1 WT*). As a consequence, *Rock1 NC* expressing MEFs did not rapidly contract or produce significant numbers of membrane blebs compared to *Rock1 WT* expressing MEFs. Not only were there morphological differences, *Rock1 NC* expressing MEFs also released more lactate dehydrogenase (LDH; a marker of cytoplasmic leakage) and greater levels of the damage-associated molecular pattern (DAMP) protein high mobility group B1 (HMGB1) (Yang et al., 2015), indicating that cell death was more necrotic-like than for MEFs expressing ROCK1 *WT*.

Given the elevated rate of apoptosis in cells expressing high levels of *c-MYC* (Evan et al., 1992) and evidence of significant apoptosis in B cell lymphoma tumours in *Eµ-Myc* mice (Park et al., 2005), we sought to determine if the aberrant contractile force generation and consequent altered apoptotic morphological changes in *Rock1 NC* cells would affect B cell lymphoma induced by *c-MYC*. Crossing *Eµ-Myc* mice with *Rock1 NC* or *Rock1 WT* mice revealed that the absence of caspase-cleavage of ROCK1 resulted in a significant 45% increase in median survival, and a shift towards an increased incidence of thymic lymphomas in *Eµ-Myc; Rock1 NC* mice The prolonged survival of *Eµ-Myc; Rock1 NC* mice was associated with decreased bone marrow cellularity and altered cell cycle progression, although there were no significant effects on B cell differentiation. Despite there being significantly fewer circulating macrophages in *Eµ-Myc; Rock1 NC* mice, there was a dramatic 18-fold increase in the proportion of macrophages in total bone marrow cells, with a significant shift towards the classical M1 pro-inflammatory state. In transplantation experiments, the survival of lethally irradiated *Rock1 WT* mice was 48% longer when receiving pre-neoplastic *Eµ-Myc; Rock1 NC* bone marrow cells relative to *Eµ-Myc; Rock1 WT* bone marrow cells, indicating that the survival differences were intrinsic properties of the transplanted bone marrow cells and not host tissues. In contrast, transplantation of established lymphoma cell lines isolated from *Eµ-Myc; Rock1 NC* or *Eµ-Myc; Rock1 WT* mice resulted in no differences in the survival of lethally irradiated *Rock1 WT* recipient mice. Taken together, the results of this study indicate that the caspase-cleavage of ROCK1 influences the characteristics of *c-MYC* induced B cell lymphomas, which was associated with greater numbers of bone marrow macrophages, altered bone marrow cell cycle profiles and decreased cellularity.

## Results

### Caspase-resistant *Rock1 NC* prolongs the survival of *Eµ-Myc* expressing mice

Primary lymphoma cell lines were established from mice expressing a single *Eµ-Myc* allele, and either two alleles of wild-type *Rock1* (*Eµ-Myc; Rock1 WT*) or two alleles of D1113A caspase-resistant *Rock1* (*Eµ-Myc; Rock1 NC*). Consistent with previous results in mouse fibroblasts that demonstrated the resistance of ROCK1 NC to caspase-mediated proteolysis (Julian et al., 2021), treatment of lymphoma cells with a combination of the BCL-2-selective inhibitor ABT199 (Souers et al., 2013) plus cycloheximide (Chx) to induce apoptosis resulted in ROCK1 WT but not ROCK1 NC protein cleavage (**Fig. 1A**). Treatment with the ROCK selective inhibitor H1152 did not affect ROCK1 WT cleavage, indicating that the proteolysis was not dependent on kinase activity (**Fig. 1A**).

**Fig. 1.**
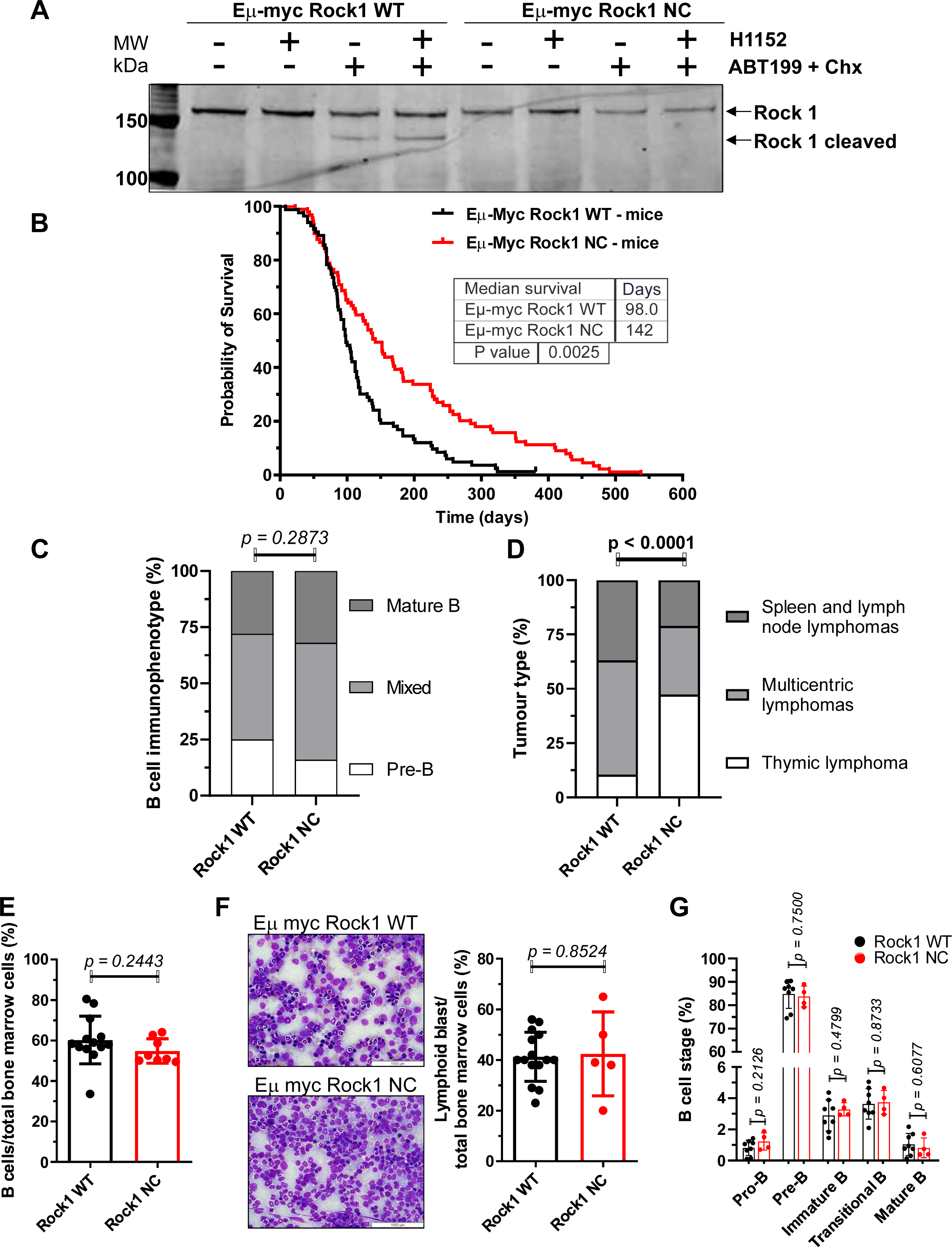
Prolonged survival of *Eµ-Myc; Rock1 NC* mice. **A.** Primary lymphoma cells from *Eµ-Myc; Rock1 WT* or *Eµ-Myc; Rock1 NC* mice were left untreated, treated with the ROCK inhibitor H1152 (10 µM, 4 h), a combination of the BCL-2-selective inhibitor ABT199 (10 µM) plus cycloheximide (Chx, 10 ng/mL, 1h) to induce apoptosis, or a combination of H1152, ABT199 and Chx. Representative western blot of cell lysates probed with anti-ROCK1 antibody revealed that the smaller ∼ 130 kDa caspase cleaved form is visible only in *Rock1 WT,* and not *Rock1 NC,* cells that had been treated with ABT199 plus CHX, indicating that the ∼ 160 kDa ROCK1 NC protein was caspase resistant as previously shown (Coleman et al., 2001; Julian et al., 2021). **B.** Kaplan-Meier estimate of the probability of survival *Eµ-Myc; Rock1 WT* (n = 82) or *Eµ-Myc; Rock1 NC* mice (n = 89). Censored mice are indicated by a vertical tick mark. *P* value was determined by Gehan-Breslow-Wilcoxon test. **C.** Immunophenotyping of lymphomas isolated from *Eµ-Myc; Rock1 WT* (n = 32) and *Eµ-Myc; Rock1 NC* (n = 25) mice by flow cytometry. *P* value was determined by Chi-square test. **D.** Incidence of tumour sites for *Eµ-Myc; Rock1 WT* (n = 19) and *Eµ-Myc; Rock1 NC* (n = 19). Thymic lymphomas were only found in thymus, lymphomas were located in spleen and lymph nodes (with no thymic enlargement), or multicentric lymphomas in which thymus, spleen and lymph nodes were all grossly affected. *P* value was determined by Chi-square test. **E.** Ratio of B cells to total bone marrow cells was determined by flow cytometry for *Eµ-Myc; Rock1 WT* (n = 13) and *Eµ-Myc; Rock1 NC* (n = 8) pre-neoplastic mice. **F.** *Left panels;* Representative photomicrographs of May-Grunwald-Giemsa-stained *Eµ-Myc; Rock1 WT* and *Eµ-Myc; Rock1 NC* bone marrow cells. Scale bars = 1000 µm. *Right panel;* Percentage of lymphoid blast cells in total bone marrow populations from *Eµ-Myc; Rock1 WT* (n = 15) and *Eµ-Myc; Rock1 NC* (n = 5) mice. **G.** Differentiation stages of B cells from the bone marrow of *Eµ-Myc; Rock1 WT* (n = 8) and *Eµ-Myc; Rock1 NC* (n = 4) mice. Graphs in **E, F, G** show means ± standard deviation, and *p* values determined by unpaired Student’s *t*-tests between the indicated groups. Data points represent individual mice at ≤ 8 weeks of age.

Mice expressing the *Eµ-Myc* transgene were examined three times per week for signs of morbidity or lymphoma emergence. *Eµ-Myc; Rock1 WT* mice succumbed to lymphoma with a median survival of 98 days and maximum survival of 324 days, while *Eµ-Myc; Rock1 NC* mice had 45% longer median survival of 142 days and maximum survival of 538 days (**Fig, 1B**). B cell immunophenotyping in mice with overt lymphomas showed no significant differences between *Eµ-Myc; Rock1 WT* and *Eµ-Myc; Rock1 NC* mice, with comparable proportions of pre-B (c-Kit^-^, B220^+^, cell surface immunoglobulin (sIg)^low^), mixed (c-Kit^-^, B220^+^, sIg^Low+High^) and mature (c-Kit^-^, B220^+^, sIg^High^) B cells (**Fig. 1C, Fig. S1**). There was a significant change in the observed disease spectrum, with an over four-fold increased proportion of mice with thymic B cell lymphomas, and concomitant ∼40% decreases in the proportions of B cell lymphomas located in spleen and lymph nodes as well as multicentric B cell lymphomas in which thymus, spleen and lymph nodes were grossly affected tumour sites (**Fig. 1D**) in *Eµ-Myc; Rock1 NC* mice relative to *Eµ-Myc; Rock1 WT* mice. Despite the altered distribution of tumour sites, there were no significant differences between the genotypes in body weights (**Fig. S2A**), or thymus (**Fig. S2B**), spleen (**Fig. S2C**) or lymph node (**Fig. S2D**) weights proportional to individual body weights. Taken together, these observations indicate that the inability to generate the hyperactive form of ROCK1 *via* caspase cleavage was associated with prolonged survival of *Eµ-Myc* expressing mice.

In healthy pre-neoplastic mice (≤ 8 week old) there were no significant differences between *Eµ-Myc; Rock1 WT* and *Eµ-Myc; Rock1 NC* mice in the proportion of B220^+^ B cells relative to total bone marrow cell numbers (**Fig. 1E, Fig. S3**). Similarly, there were no significant differences in the proportions of May-Grunwald-Giemsa positive stained lymphoid blast cells between the genotypes (**Fig. 1F**). Nor were there differences in B cell differentiation, with no significant differences from pro-B to mature B cell stages (**Fig. 1G, Fig. S3**). These observations indicate that the prolonged survival of *Eµ-Myc* expressing mice bearing the *Rock1 NC* mutation was not associated with effects on B cell differentiation state.

### Reduced bone marrow cellularity and altered cell cycle profiles in *Eµ-Myc; Rock1 NC* mice

Cross sections of haematoxylin-stained femurs from healthy pre-neoplastic (≤ 8 week old) mice revealed visibly lower bone marrow cell densities in *Eµ-Myc; Rock1 NC* mice relative to *Eµ-Myc; Rock1 WT* mice (**Fig. 2A**, left and right panels). Counting total bone marrow cells from individual femurs indicated that there were significantly ∼60% fewer cells in *Eµ-Myc; Rock1 NC* femurs compared to *Eµ-Myc; Rock1 WT* femurs (**Fig. 2B**). To determine if the differences in total cell number were associated with altered proliferation, mice were injected with 250 µl BrdU (10 mg/mL) intraperitoneally 2 hours prior to sacrifice, then bone marrow cells were isolated, fixed, and stained for BrdU incorporation and DNA content with propidium iodide prior to analysis by flow cytometry. *Eµ-Myc; Rock1 NC* mice were found to have a significantly >50% greater proportion of cells in the G_0_/G_1_ cell cycle phase, 23% lower in the DNA synthetic S phase, and 88% lower in the apoptotic sub-G_1_ phase relative to *Eµ-Myc; Rock1 WT* mice (**Fig. 2C**). There were no significant differences in the proportion of cells in the G_2_/M cell cycle phase between the genotypes (**Fig. 2C**). These results indicate that the reduced bone marrow cell numbers in *Eµ-Myc; Rock1 NC* femurs was associated with cell cycle changes associated with reduced cell proliferation.

**Fig. 2.**
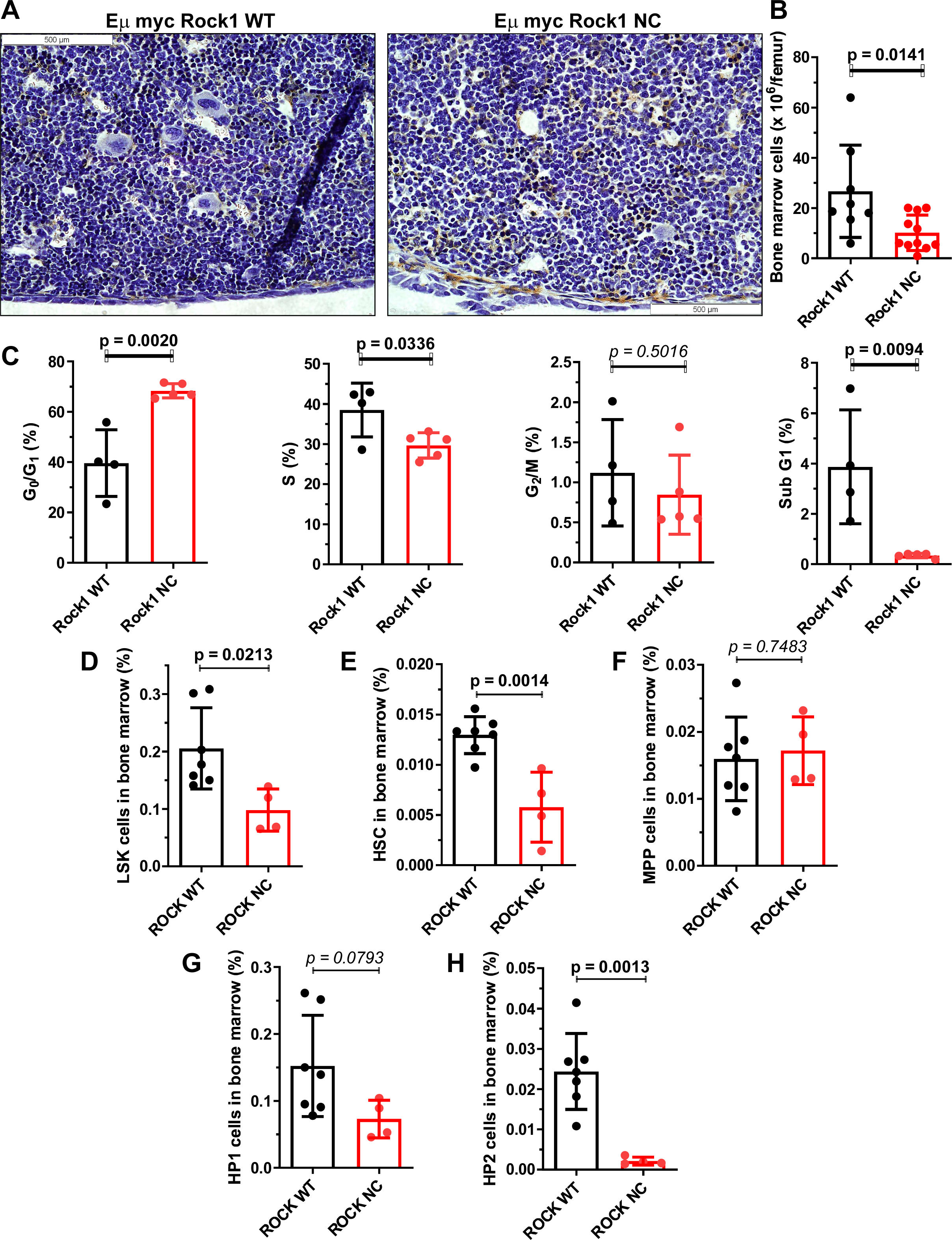
Reduced cellularity and proliferation in pre-neoplastic *Eµ-Myc; Rock1 NC* bone marrow does not affect B cell proportions or differentiation. **A.** Cross section of haematoxylin-stained femur and bone marrow from *Eµ-Myc; Rock1 WT* and *Eµ-Myc; Rock1 NC* mice. Scale bars = 500 µm. **B.** Number of total bone marrow cells per femur from *Eµ-Myc; Rock1 WT* (n = 8) and *Eµ-Myc; Rock1 NC* (n = 11) mice. **C.** Cell cycle profiles determined after *Eµ-Myc; Rock1 WT* (n = 4) and *Eµ-Myc; Rock1 NC* (n = 5) mice were injected with 250 µl BrdU (10 mg/mL) intraperitoneally 2 hours prior to sacrifice. BrdU incorporation and DNA content (propidium iodide) were determined by flow cytometry. Percentages of total bone marrow cells of: **D.** Lineage marker (Lin; CD3, CD4, CD8, CD11b, B220, GR-1, TER119) negative Sca1^+^ c-Kit^+^ (LSK) hematopoietic stem cells (HSC). **E.** Lin^-^ CD48^low^ CD150^high^ HSCs. **F.** Lin^-^ CD48^low^ CD150^low^ multi-potent progenitor (MPP) cells **G.** Lin^-^ CD48^high^ CD150^low^ hematopoietic progenitor 1 (HP1) cells. **H.** Lin^-^ CD48^high^ CD150^high^ HP2 cells; for *Eµ-Myc Rock1 WT* (n = 7) and *Eµ-Myc Rock1 NC* (n = 4) mice. All *p* values were determined by unpaired Student’s *t*-tests between indicated groups. All graphs show means ± standard deviation, with data points representing individual mice ≤ 8 weeks of age.

Ectopic *c-MYC* over-expression was previously found to reduce hematopoietic stem cell populations due a lower capacity for self-renewal resulting from the increased proliferation and differentiation of progenitor cells (Wilson et al., 2004). Given these previous observations, there was a possibility that there would be differences in hematopoietic stem cell numbers and differentiation states. There were significantly fewer lineage marker (Lin; CD3, CD4, CD8, CD11b, B220, GR-1, TER119) negative Sca1^+^ cKit^+^ (LSK) hematopoietic stem cells (HSC) in *Eµ-Myc; Rock1 NC* mice relative to *Eµ-Myc; Rock1 WT* mice as a proportion of total bone marrow cells (**Fig. 2D, Fig. S3**). Further characterization of the LSK cells revealed that there were significantly fewer Lin^-^ CD48^low^ CD150^high^ HSC (**Fig. 2E**) in *Eµ-Myc; Rock1 NC* bone marrows relative to *Eµ-Myc; Rock1 WT* bone marrows. There were no differences in Lin^-^ CD48^low^ CD150^low^ multi-potent progenitor (MPP) cells (**Fig. 2F**) or Lin^-^ CD48^high^ CD150^low^ hematopoietic progenitor 1 (HP1) cells (**Fig. 2G**). However, there were significantly fewer Lin^-^ CD48^high^ CD150^high^ HP2 cells in in *Eµ-Myc; Rock1 NC* mice relative to *Eµ-Myc; Rock1 WT* mice (**Fig. 2H**). Taken together, these results are consistent with *c-MYC* expression combined with the altered cell cycle profiles in the bone marrows of *Eµ-Myc; Rock1 NC* mice having a pronounced effect on reducing the numbers of hematopoietic stem cells and hematopoietic progenitor 2 cells.

Given the observed effects on reducing the proportion of hematopoietic stem cells in *Eµ-Myc; Rock1 NC* bone marrows, the levels of circulating blood cells were examined. Clinical pathology hematological analysis revealed no significant differences between pre-neoplastic *Eµ-Myc; Rock1 WT* and *Eµ-Myc; Rock1 NC* mice in the levels of circulating red blood cell numbers (**Fig. 3A**), hemoglobin (**Fig. S4A**), hematocrit (**Fig. S4B**), mean corpuscular volume (**Fig. S4C**), mean corpuscular hemoglobin (**Fig. S4D**), mean corpuscular hemoglobin concentration (**Fig. S4E**) or red cell distribution width (**Fig. S4F**). Qualitative hematological analysis also revealed no significant differences in polychromasia, red blood cell spiculing, hemoglobin crystals, or the presence of large lymphoid or pyknotic cells (**Table S1**). There were also no significant differences in platelet numbers (**Fig. 3B**), mean platelet volume (**Fig. S4G**), plateletcrit (**Fig. S4H**), or platelet distribution width (**Fig. S4I**). In addition, there were no significant differences in total neutrophil (**Fig. 3C**), eosinophil (**Fig. 3D**), monocyte (**Fig. 3E**) or lymphocyte numbers (**Fig. 3F**). More detailed cell classification by flow cytometry (**Fig. S5**) revealed that *Eµ-Myc; Rock1 NC* mice had no significant differences in B220^+^ B cell numbers relative to *Eµ-Myc; Rock1 WT* mice (**Fig 4A**) but did have ∼80% lower levels of CD11c^+^ dendritic cells (DC, **Fig. 4B**) and 73% lower levels of Lin^-^ Ly6C^low^ CD115^+^ F4/80^+^ macrophages (**Fig. 4C**) than in *Eµ-Myc; Rock1 WT* mice. Similarly, although clinical pathological analysis revealed no significant differences in total lymphocyte numbers (**Fig. 3F**), more detailed flow cytometric analysis (**Fig. S6**) indicated that there were significantly 65% lower levels of CD3^+^ T cells (**Fig. 4F**) and CD4^+^ CD69^+^ effector T cells (**Fig. 4E**), as well as ∼90% lower levels of CD4^-^ CD8α^+^ cytotoxic T lymphocytes (CTL; **Fig. 4F**), CD4^+^ CD25^+^ regulatory T cells (Treg; **Fig. 4G**) and CD3^-^ CD49b^+^ natural killer cells (NK; **Fig. 4H**) in *Eµ-Myc; Rock1 NC* mice than in *Eµ-Myc; Rock1 WT* mice. Therefore, although the significantly reduced bone marrow cellularity and proportionately lower HSC levels in *Eµ-Myc; Rock1 NC* mice were not associated with lower levels of major types of blood cells, there were effects on the levels of specific sub-types of monocytes and lymphocytes indicating that differentiation pathways were likely affected.

**Fig. 3.**
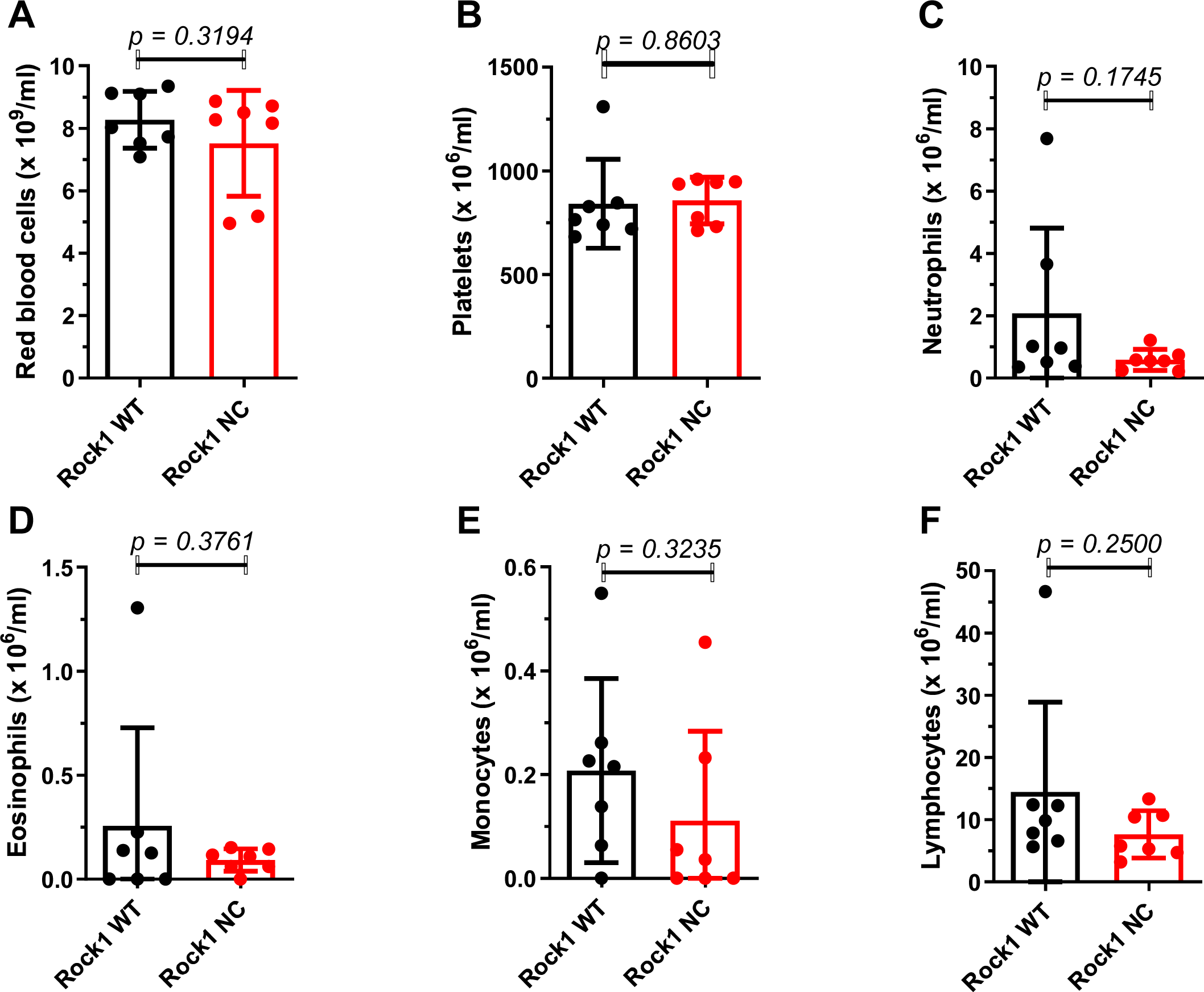
No differences in the circulating levels of major blood cell typess in pre- neoplastic *Eµ-Myc; Rock1 NC* mice. Haematological profiling of pre-neoplastic *Eµ-Myc; Rock1 WT* (n = 7) and *Eµ-Myc; Rock1 NC* (n = 7) mice for the number of circulating **A.** red blood cells, **B.** platelets. **C.** neutrophils, **D.** eosinophils, **E.** ^monocytes^ and **F.** lymphocytes.

**Fig. 4.**
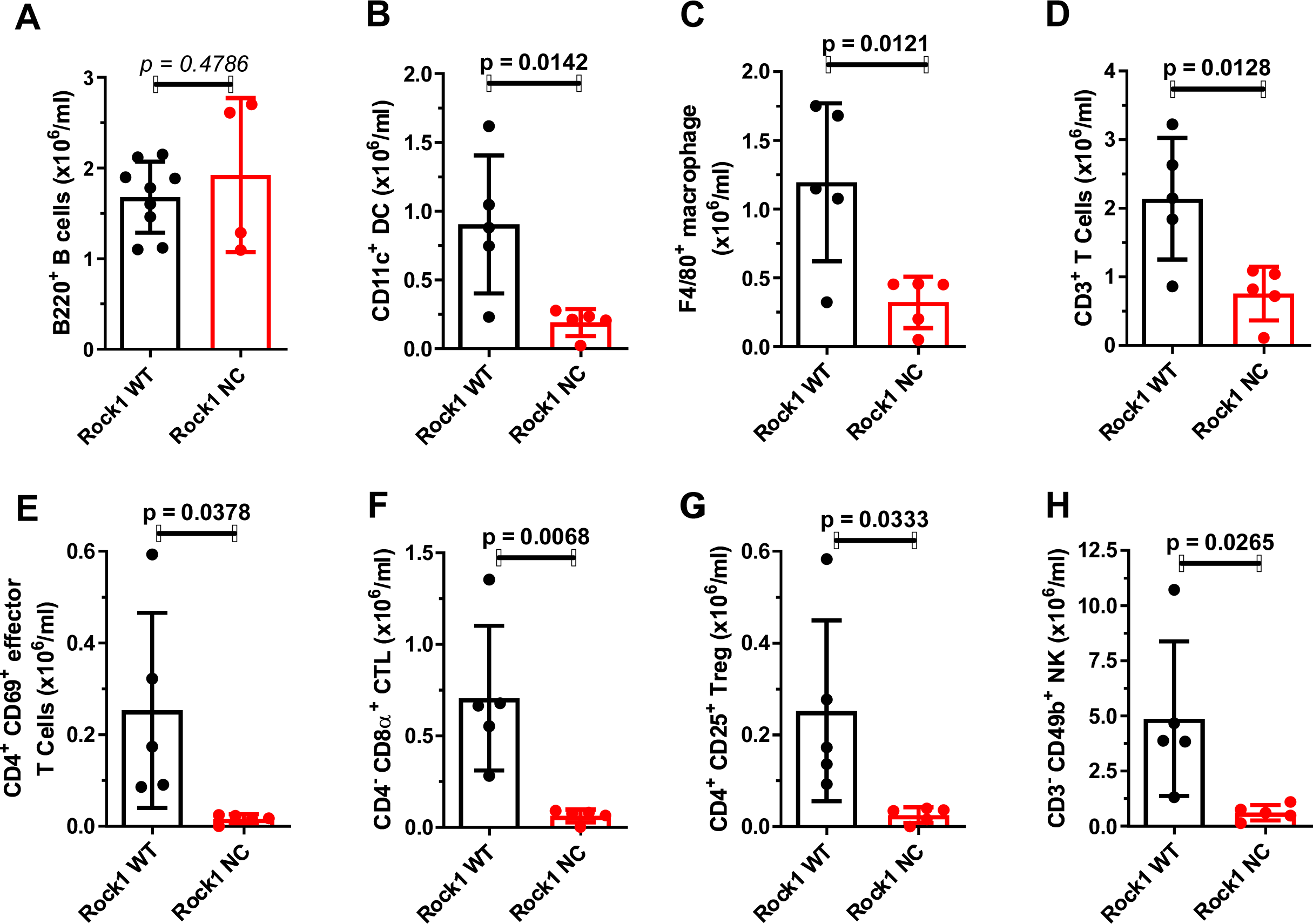
Lower levels of circulating lymphocyte sub-types in *Eµ-Myc; Rock1 NC* mice. Flow cytometry analysis of *Eµ-Myc; Rock1 WT* (n = 5) and *Eµ-Myc; Rock1 NC* (n = 5) mice for the number of circulating **A.**B220^+^ B cells, **B.** CD11c^+^ dendritic cells (DC), **C.** F4/80^+^ macrophages, **D.** CD3^+^ T cells, **E.** CD4^+^ CD69^+^ effector T cells, **F.** CD4^+^ CD8α^+^ cytotoxic T lymphocytes (CTL), **G.** CD4^+^ CD25^+^ regulatory T cells (Treg) and **H.** CD3^-^ CD49b^+^ natural killer (NK) cells. All *p* values were determined by unpaired Student’s *t*-tests between indicated groups. All graphs show means ± standard deviation, with data points representing individual mice ≤ 8 weeks of age.

### Bone thickening in pre-neoplastic *Eµ-Myc; Rock1 NC* mice

When the tibias of age-matched pre-neoplastic *Eµ-Myc; Rock1 WT* and *Eµ-Myc; Rock1 NC* mice were scanned with a Bruker micro computed tomography (micro-CT) Skyscan1172 and compared using two dimensional (2D) scans (**Fig. 5A**, top panels) or three dimensional (3D) reconstructions (**Fig. 5A**, bottom panels), it was evident that the trabecular bones were thicker in the *Eµ-Myc; Rock1 NC* mice. Quantitative micro-CT analysis from 2D and 3D measurements revealed that *Eµ-Myc; Rock1 NC* mice had significantly 10% greater trabecular thickness (**Fig. 5B**), 19% greater intersection surfaces (**Fig. 5C**), 9% greater structure model index (a measurement of 3D structures in terms of their composition by plates and rods as described in Hildebrand and Rüegsegger (Hildebrand and Rüegsegger, 1997) (**Fig. 5D**), 11% greater tissue surface (tissue indicating the object identified in the 3D volume of interest, **Fig. 5E**) and 17% greater tissue volume (**Fig. 5F**). There were no significant differences in bone volume, bone/tissue volume ratio, bone surface density, bone surface, bone surface/volume ratio, trabecular separation, trabecular number, trabecular pattern factor, or X, Y or Z centroids (**Fig. S7A-K**).

**Fig. 5.**
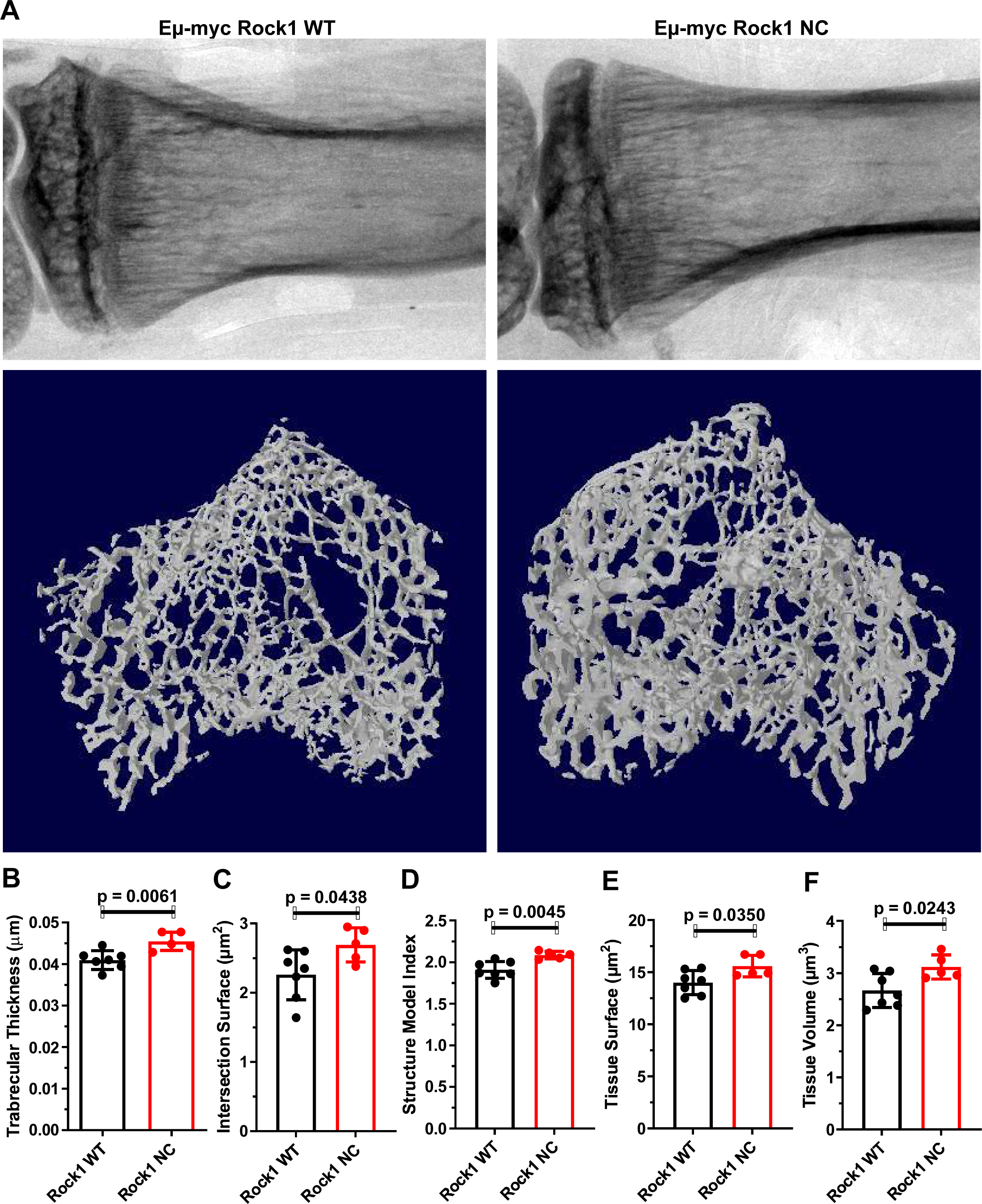
Thicker tibia trabecular bones in pre-neoplastic *Eµ-Myc; Rock1 NC* mice. **A.** Representative two-dimensional scans (upper panels) and three-dimensional reconstructions (lower panels) of tibia by micro computed tomography (micro-CT) from *Eµ-Myc; Rock1 WT* and *Eµ-Myc; Rock1 NC* mice. **B.** Trabecular thickness, **C.** intersection surface, **D.** structure model index, **E.** tissue surface and **F.** tissue volume determined from micro-CT scans of tibia from *Eµ-Myc; Rock1 WT* (n = 7) and *Eµ-Myc; Rock1 NC* (n = 5) mice. All *p* values were determined by unpaired Student’s *t*-tests between indicated groups. All graphs show means ± standard deviation, with data points representing individual mice ≤ 8 weeks of age.

### Elevated macrophage numbers in *Eµ-Myc; Rock1 NC* bone marrow are not due to altered macrophage properties

In addition to promoting cell proliferation, *c-MYC* has pro-apoptotic effects mediated via the p19^ARF^–MDM2–p53 pathway (Eischen et al., 1999). Notably, the pro-apoptotic actions of the *Eµ-Myc* transgene were previously associated with increased numbers of bone marrow macrophages relative to non-transgenic control mice (Jacobsen et al., 1994). Since we previously demonstrated that the *Rock1 NC* mutation reduced contractile force generation during apoptosis that resulted in a necrotic-like form of cell death associated with increased inflammation (Julian et al., 2021; Naylor et al., 2022), the effects on macrophage numbers and polarization states were examined in *Eµ-Myc; Rock1 WT* and *Eµ-Myc; Rock1 NC* mice by flow cytometry. Despite there being fewer circulating macrophages in pre-neoplastic *Eµ-Myc; Rock1 NC* mice (**Fig. 4C**), there was a significant 18-fold greater number of macrophages in the bone marrows of *Eµ-Myc; Rock1 NC* mice relative to *Eµ-Myc; Rock1 WT* mice (**Fig. 6A**), with a significant shift towards the classically defined pro-inflammatory MHC-II^high^ CD206^low^ M1 state (**Fig. 6B**) and away from the anti-inflammatory MHC-II^low^ CD206^high^ M2 state (**Fig. 6C**). This definition of macrophage polarization comes with the caveat that there is a lack of precisely defined and universally accepted criteria to score phenotypes. Consistent with these observations, there were also significantly > 50% more macrophages in *Rock1 NC* thymi relative to *Rock1 WT* thymi at ≤ 8 weeks of age in the absence of the *Eµ-Myc* transgene, suggesting that spontaneous thymocyte apoptosis similarly evoked greater macrophage recruitment (**Fig. S8**). To determine if there were differences in the intrinsic properties of macrophages that might contribute to their relative abundance in *Eµ-Myc; Rock1 NC* bone marrows, primary macrophages were isolated from *Eµ-Myc; Rock1 WT* or *Eµ-Myc; Rock1 NC* mice and examined *ex vivo* (**Fig. S9A**). Tracking random macrophage migration (**Fig. S9B,C**) revealed no significant differences in mean accumulated distances travelled over 24 hours (**Fig. 6D**), mean Euclidean distances travelled (**Fig. 6E**), mean cell velocities (**Fig. 6F**) or migration directionality (**Fig. 5G**). When the ability of macrophages to take up fluorescent latex beads was assessed, there were no differences in the proportion of macrophages that had ingested beads or in the number of beads ingested per cell (**Fig. 6H, Fig. S10**). Furthermore, the difference in macrophage numbers in *Eµ-Myc; Rock1 NC* bone marrows was not replicated in an acute peritonitis model in which zymosan, a polysaccharide cell wall component derived from *Saccharomyces cerevisiae* (Cash et al., 2009), was injected into the peritoneal cavities of *Eµ-Myc; Rock1 WT* or *Eµ-Myc; Rock1 NC* mice to induce acute inflammation. At 24 hours following zymosan injection, the numbers of peritoneal white blood cells (**Fig. 6I**) or F4/80^+^ macrophages, either as total numbers (**Fig. 6J**) or as the proportion of white blood cells (**Fig. 6K**), were equivalent in *Eµ-Myc; Rock1 WT* and *Eµ-Myc; Rock1 NC* mice. Taken together, these results indicate that there were no detectable differences in the behaviours or properties of macrophages between *Eµ-Myc; Rock1 WT* or *Eµ-Myc; Rock1 NC* mice that would likely contribute to the elevated presence of macrophages in the bone marrows of *Eµ-Myc; Rock1 NC* mice (**Fig. 6A**).

**Fig. 6.**
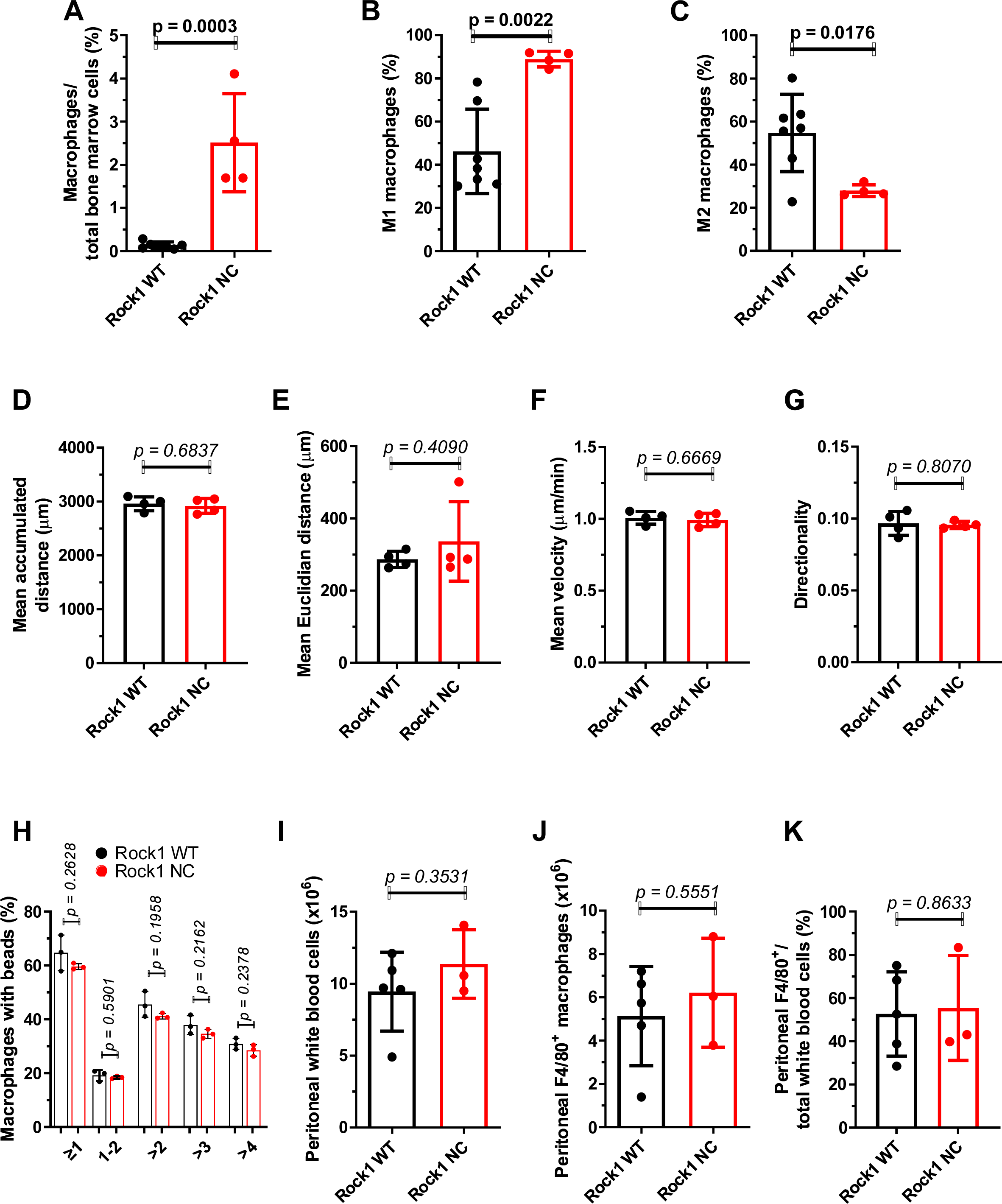
Greater macrophage numbers in pre-neoplastic *Eµ-Myc; Rock1 NC* bone marrow. **A.** Percentage macrophages in the bone marrows of pre-neoplastic *Eµ-Myc; Rock1 WT* (n = 7) and *Eµ-Myc; Rock1 NC* (n = 4) mice was determined by flow cytometry. The percentages of macrophages that were polarized as **B.** M1 or **C.** M2 were determined by flow cytometry. The motility of primary peritoneal macrophages was tracked over 24 hours, and quantified as the **D.** mean accumulated distance traveled, **E.** mean Euclidean distance traveled, **F.** mean velocity and **G.** directionality (the ratio of Euclidean to accumulated distances) for *Eµ-Myc; Rock1 WT* (n = 4) and *Eµ-Myc; Rock1 NC* (n = 4) mice. Peritoneal macrophages isolated from each mouse were plated in triplicate wells, each well was imaged in three positions to give mean values from each well, which were then averaged to give results for each mouse. Mean number of tracked cells per well position = 87. **H.** Bone marrow derived macrophages from *Eµ-Myc; Rock1 WT* (n = 3) and *Eµ-Myc; Rock1 NC* (n = 3) mice were incubated for 2 hours with serum-opsonized 2 µm diameter FluoSpheres, then the number of beads per cell determined by flow cytometry. In a model of acute peritonitis in which *Eµ-Myc; Rock1 WT* (n = 5) and *Eµ-Myc; Rock1 NC* (n = 3) mice received intraperitoneal injections of 1 mg Zymosan, 24 hours later the numbers of **I.** white blood cells and **J.** F4/80^+^ macrophages in the peritoneal cavity were analyzed by flow cytometry. **K.** The ratio of F4/80^+^ macrophages to total white blood cells. All *p* values were determined by unpaired Student’s *t*-tests between indicated groups. All graphs show means ± standard deviation, with data points representing individual mice ≤ 8 weeks of age.

### Transplantation of pre-neoplastic *Eµ-Myc; Rock1 NC* bone marrow into *Rock1 WT* hosts recapitulates prolonged survival

To determine if the differences in survival between *Eµ-Myc; Rock1 WT* and *Eµ-Myc; Rock1 NC* mice were intrinsic to bone marrow cells or if there were additional extrinsic contributory factors, equal numbers of bone marrow cells from the two genotype *CD45.2* donors were transplanted into irradiated *CD45.1 Rock1 WT* host mice (Latif et al., 2021). Similar to the enhanced survival of *Eµ-Myc; Rock1 NC* genetically modified mice relative to *Eµ-Myc; Rock1 WT* (**Fig. 1B**), *Rock1 WT* mice receiving *Eµ-Myc; Rock1 NC* bone marrow cells lived significantly 48% longer (median survival 37 days, maximum survival 52 days) than mice receiving *Eµ-Myc; Rock1 WT* bone marrow cells (median survival 25 days, maximum survival 28 days) (**Fig. 7A**). The body weights of mice that received either bone marrow genotype were not significantly different at experimental endpoints (**Fig. S11A**), nor was the ratio of CD45.2^+^/total white blood cells (**Fig. S11B**). In addition, the ratio of CD45.2^+^ B cells/total white blood cells was not different between the mice receiving *Eµ-Myc; Rock1 WT* or *Eµ-Myc; Rock1 NC* bone marrow transplants (**Fig. S11C**). Taken together, these observations indicate that the engraftment efficiency of the transplanted bone marrows was comparable between *Eµ-Myc; Rock1 WT* and *Eµ-Myc; Rock1 NC* donors. Therefore, the prolonged survival of mice with transplanted *Eµ-Myc; Rock1 NC* cells can be attributed to phenotypic differences resulting from the resistance of mutant Rock1 to caspase cleavage and consequent hyperactivation during apoptotic cell death.

**Fig. 7.**
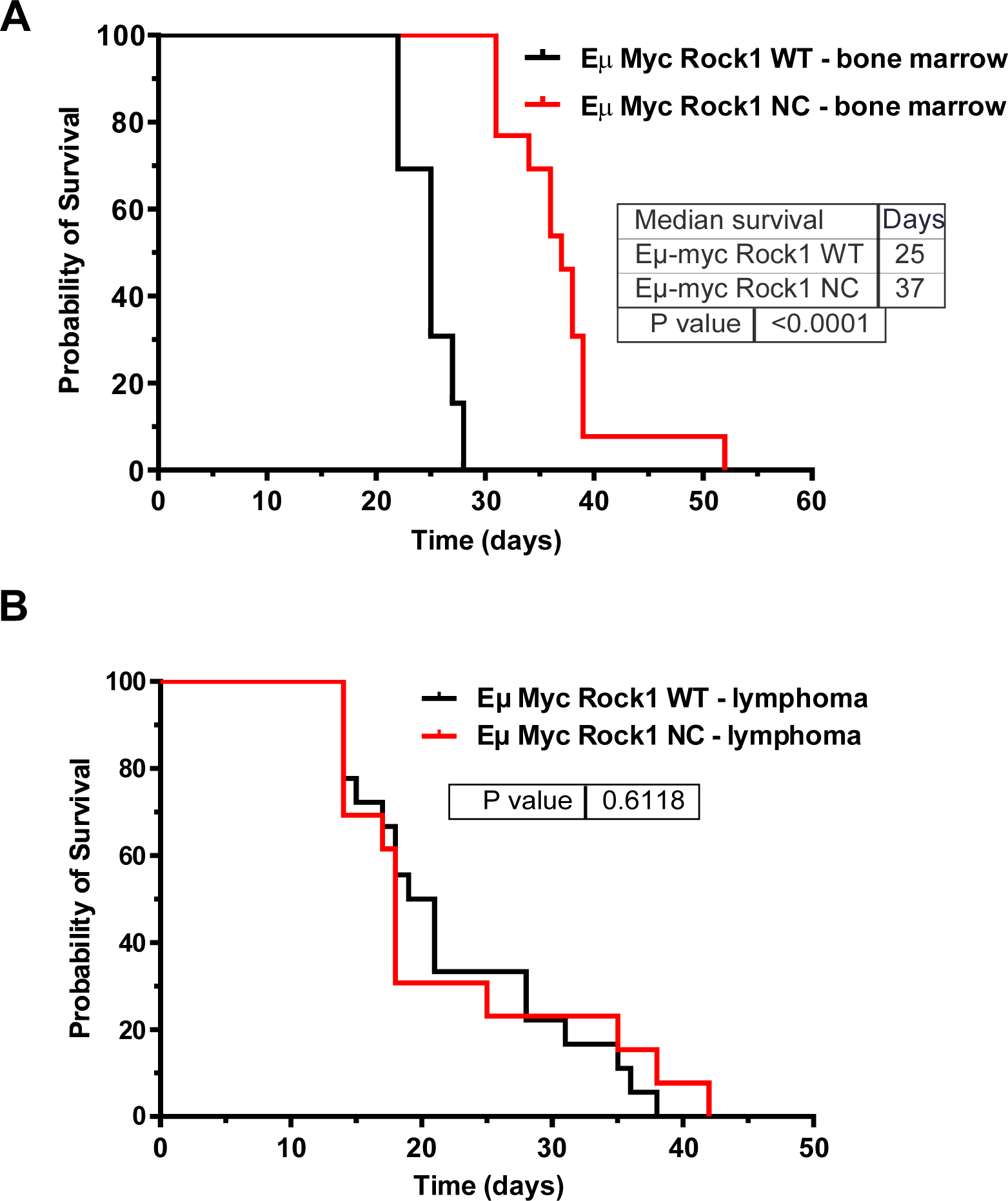
Transplanted bone marrow from *Eµ-Myc; Rock1 NC* provides a survival advantage. **A.** Kaplan-Meier estimate of the probability of survival of irradiated *Rock1 WT* mice transplanted with bone marrow cells from *Eµ-Myc; Rock1 WT* (n = 13) or *Eµ-Myc; Rock1 NC* (n = 13) mice. *P* value was determined by Gehan-Breslow-Wilcoxon test. **B.** Kaplan-Meier estimate of the probability of survival of irradiated *Rock1 WT* mice transplanted with lymphoma cells from *Eµ-Myc; Rock1 WT* (n = 18) or *Eµ-Myc; Rock1 NC* (n = 13) mice. *P* value was determined by Gehan-Breslow-Wilcoxon test.

To determine if the survival advantage conferred by the *Rock1 NC* mutation was dependent on the transformation status of lymphoma cells, irradiated *Rock1 WT* recipient mice were transplanted with established primary lymphoma cells isolated from *Eµ-Myc; Rock1 WT* or *Eµ-Myc; Rock1 NC* mice that had displayed no significant differences in growth rates *in vitro* (**Fig. S11D**). In contrast to the significantly longer survival of mice receiving pre-neoplastic *Eµ-Myc; Rock1 NC* bone marrow (**Fig. 7A**), there was no difference in the survival of mice transplanted with *Eµ-Myc; Rock1 WT* (median survival 20 days, maximum survival 42 days) or *Eµ-Myc; Rock1 NC* lymphoma cells (median survival 18 days, maximum survival 38 days) (**Fig. 7B**). Therefore, additional mutations acquired during transformation likely made the lymphoma cells independent of microenvironmental responses associated with expression of the caspase-resistant ROCK1 NC protein in the transplanted bone marrows

## Discussion

The AlphaFold (Jumper et al., 2021) predicted structure of full-length mouse ROCK1 (**Video** 1: AF-P70335-F1) places Asp1113 in the caspase cleavage site (**Fig. 8**; red) on a relatively unstructured loop that connects the final α helix (**Fig. 8**; pink) to the globular pleckstrin homology (PH) and cysteine-rich (CR) regions (**Fig. 8**; dark blue and light blue, respectively). It is unclear how removal of the PH and CR domains (**Fig. 8B**) would result in activation of the kinase domain (**Fig. 8**; light green); one possibility is that it leads to repositioning of the coiled-coils to enable adoption of an active conformation and/or greater accessibility to protein substrates. The Rho-binding domain (**Fig. 8**; dark green) is also positioned away from the kinase domain at the end of the penultimate α helix. Nevertheless, binding of GTP-loaded active RhoA has been shown to increase ROCK1 catalytic activity (Amano et al., 1997; Leung et al., 1996), possibly mediated by a similar re-positioning of the coiled-coils to facilitate kinase activity. Consistent with inhibitory effects of the coiled-coil region, early research demonstrated that progressive deletion of the carboxyl-terminal regions resulted in increasing kinase activity (Ishizaki et al., 1997; Leung et al., 1996), similar to the observed effect of caspase-mediated cleavage and removal of the carboxyl-terminal domains resulting in kinase hyperactivation.

**Fig. 8.**
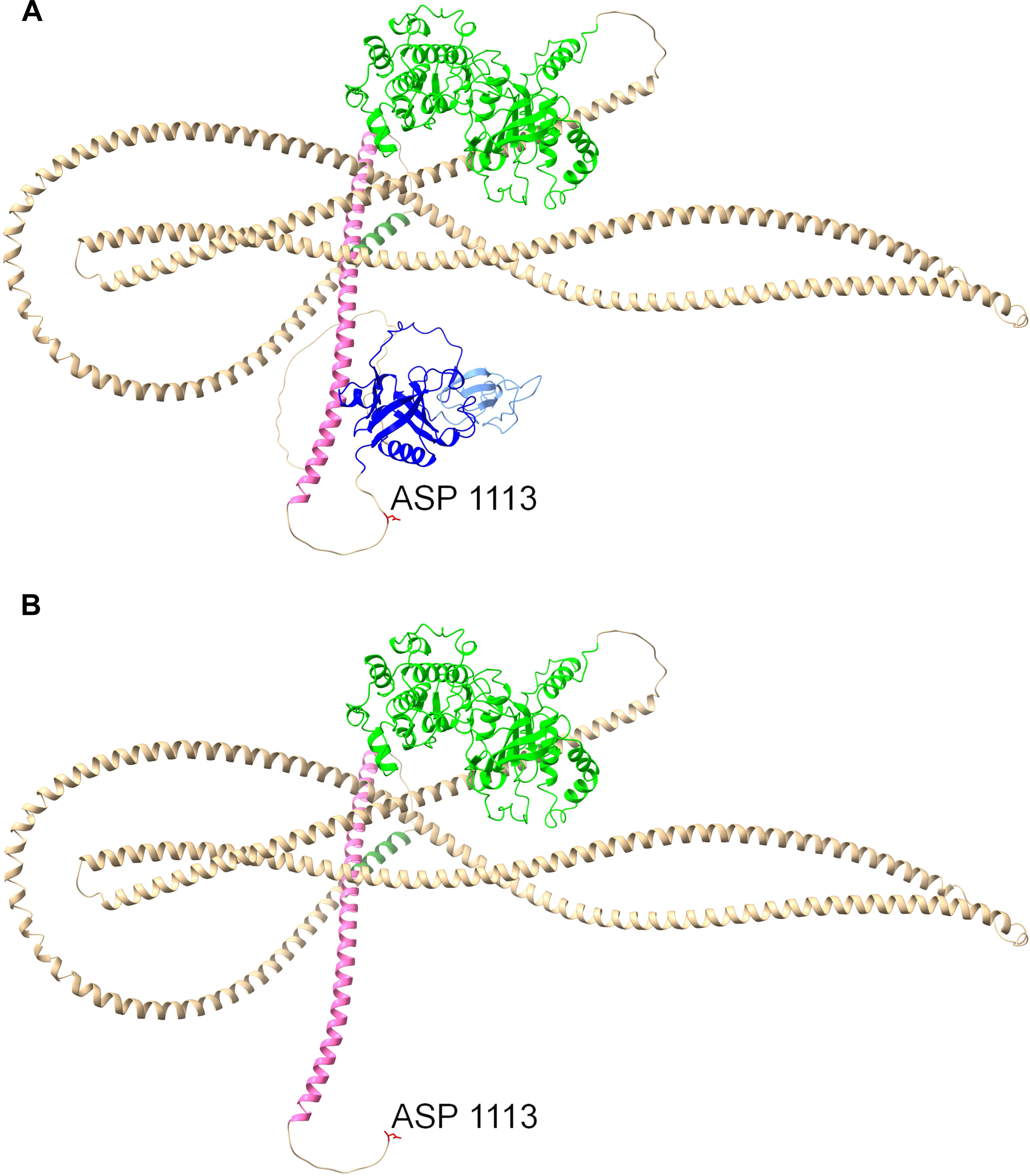
AlphaFold predicted ROCK1 structure. **A.** AlphaFold [20] predicted structure of full-length mouse ROCK1 (AF-P70335-F1). An unstructured loop connects the final α helix (pink) to the globular pleckstrin homology (PH; dark blue) and cysteine-rich (CR; light blue) regions. Kinase domain (light green) and Rho-binding domain (dark green) are separated by α helices (light brown). Aspartic acid (ASP) 1113 is the final amino acid of the caspase recognition sequence. **B.** The proteolytic cleavage of ROCK1 by activated caspases removed the terminal PH and CR domains.

The *Rock2* gene is the primordial homologue, more closely related to the isoform found in more primitive organisms. The *Rock1* gene likely is a product of the whole genome duplication event that occurred approximately 600 million years ago which enabled the evolution of complex vertebrates (Blomme et al., 2006). Given the absolute conservation of the *Rock1* caspase cleavage site in mammals, birds, fish and reptiles, the genetic divergence from *Rock2* that resulted in the caspase cleavage site specifically in *Rock1* must have been an early event after gene duplication.

The ubiquitous conservation of the ROCK1 caspase cleavage site and high homology of adjacent amino acids across mammalian, bird, fish, amphibian and reptile species is strongly suggestive of functional importance (Julian et al., 2021). And yet, despite the fact that mutating Asp1113 to an alanine residue rendered ROCK1 completely resistant to caspase-mediated cleavage, there were no observable effects in unchallenged homozygous *Rock1 NC* mutant mice (Julian et al., 2021). However, when subjected to a strong apoptotic stimulus, such as diethylnitrosamine (DEN) treatment (Julian et al., 2021) or tissue-selective *c-MYC* over-expression, there were adverse consequences associated with the absence of ROCK1 cleavage and kinase hyperactivation. In the case of DEN administration, there was relatively greater induction of cytokines and chemokines, indicative of a pronounced inflammatory response, and increased liver damage in *Rock1 NC* mice relative to *Rock1 WT* mice (Julian et al., 2021). In *Eµ-Myc* mice, the *Rock1 NC* mutation resulted in altered bone marrow cell cycling and lowered cell numbers, bone thickening, and reduction of some circulating monocyte and lymphocyte subtypes (**Figs. 2-4**). These results reinforce the conclusion that it is important for apoptotic cells to cleave and activate ROCK1 in order to undergo normal morphological changes for the maintenance of tissue homeostasis when encountering strong apoptotic stimuli.

Interestingly, in both DEN-treated and *Eµ-Myc* transgenic mice, the *Rock1 NC* mutation had apparently beneficial anti-cancer effects. DEN-treated *Rock1 NC* mice had fewer HCC tumours than *Rock1 WT* mice (Julian et al., 2021), and *Eµ-Myc; Rock1 NC* mice lived significantly longer than *Eµ-Myc; Rock1 WT* mice (**Fig. 1B**). The neutrophil-mediated damage amplification in *Rock1 NC* mice following DEN administration had the effect of eliminating potential tumour initiating cells; by reducing neutrophil recruitment immediately after DEN treatment there were increased HCC tumour numbers in the long-term (Julian et al., 2021). The prolonged survival of *Eµ-Myc; Rock1 NC* mice (**Fig. 1B**) is likely the result of the altered cell cycling and reduced number of cells in the bone marrow (**Fig. 2B**), possibly due to the loss of cellular contents that acted as damage associated molecular patterns (DAMP) from apoptotic cells, in a manner similar to the greater release of LDH and HMGB1 from apoptotic *Rock1 NC* MEFs relative to *Rock1 WT* MEFs (Julian et al., 2021). Additionally, the greater number and M1 polarization of macrophages in *Eµ-Myc; Rock1 NC* bone marrows (**Figs. 6A-C**) might have anti-proliferative effects, possibly due to the release of factors such as tumour growth factor β (TGFβ) (Fadok et al., 1998).

Although there are typically numerous resident bone marrow macrophages, additional circulating macrophages may be recruited in response to pro-inflammatory signals (Bozec and Soulat, 2017), which might contribute to the observed increase in bone marrow macrophages (**Fig. 6A**) and concomitant decrease in circulating macrophages (**Fig. 4C** in *Eµ-Myc; Rock1 NC* mice relative to of *Eµ-Myc; Rock1 WT* mice. An additional possibility is that the increased number of bone marrow macrophages result from a bias of macrophage specialization towards, and/or expansion of, a resident F4/80^+^ macrophage subtype called osteomacs that contribute to the maintenance of tissue homeostasis through the efficient phagocytosis of apoptotic cell debris, a process called efferocytosis (Bozec and Soulat, 2017). Osteomacs also provide pro-anabolic support to osteoblasts (Batoon et al., 2019) that promote bone formation and maintenance through the formation and deposition of mineralized bone matrix (Chen et al., 2020), which might account for the observed changes in bone characteristics, including thickening (**Fig. 5**), in *Eµ-Myc; Rock1 NC* mice relative to *Eµ-Myc; Rock1 WT* mice. In addition, if bone marrow resident macrophage specialization were biased towards the osteomac subtype, this could also result in a depletion of hematopoietic stem cell niche macrophages that normally maintain HSCs (Seyfried et al., 2020), which could be a contributory factor to the observed reduction in bone marrow HSCs (**Figs. 2D-E)** and hematopoietic progenitor 2 cells (**Fig. 2H**) in *Eµ-Myc; Rock1 NC* mice relative to *Eµ-Myc; Rock1 WT* mice. Given that there were no significant differences in red blood cell numbers between the genotypes (**Fig. 3A**), there may have been no effect on the specialization of macrophages into the erythroid island niche subtype (Li et al., 2020)

In transplantation experiments in which equal numbers of pre-neoplastic donor *Eµ-Myc; Rock1 NC* and *Eµ-Myc; Rock1 WT* cells were transplanted into *Rock1 WT* host mice, those receiving *Eµ-Myc; Rock1 NC* bone marrow cells still survived longer than mice transplanted with *Eµ-Myc; Rock1 WT* bone marrow cells (**Fig. 7A**), indicating that effects of the *Rock1 NC* mutation in tissues outside of the bone marrow compartment could not account for the relative survival advantage of these mice (**Fig. 1B**). In addition, the absence of survival advantage in mice transplanted with transformed *Eµ-Myc; Rock1 NC* lymphoma cells (**Fig. 7B**) suggests that they had become insensitive to growth inhibitory stimuli that they might be exposed to in the *Eµ-Myc; Rock1 WT* host bone marrows. These results, as well as the *in vitro* growth rates of *Eµ-Myc; Rock1 WT and Eµ-Myc; Rock1 NC* lymphoma cells (**Fig. S11D**), indicate that the *Rock1 NC* mutation itself is unlikely to have a direct effect on cell proliferation.

It might seem counterintuitive that mice with wild-type *Rock1* would fare less well in the HCC and B cell lymphoma cancer models than those with the *Rock1 NC* mutation, since the expectation is that evolution selects for the most advantageous properties. A possible explanation is that acute benefits on tissue homeostasis and function associated with apoptotic cell death producing typical non-inflammatory morphological changes when ROCK1 is activated by caspase cleavage are more advantageous than the apparent long-term disadvantage of worse cancer outcomes. When considering that cancers are most often manifested at advanced ages, generally past the age of reproduction, there would be a selective advantage for any attributes that promote a healthy condition up to and including reproductive adulthood, even if the same attributes could contribute to shorter overall lifespans in the aged. As a result, there could be positive health benefits that would contribute to positive selection for ROCK1 to be caspase cleaved to enable cell death to be less necrotic-like and inflammatory, which consequently might actually lead to worse cancer outcomes.

## Materials and methods

### Mouse models

*Rock1 NC* mice (Rock1^tm1.1Bicrq^) were generated at the CRUK Beatson Institute (BICR) as described in Julian *et al*. (Julian et al., 2021), and were bred into the C57Bl/6J background. *Eµ-Myc* mice (Tg(IghMyc)22Bri) in the C57Bl/6J background were from the Jackson Laboratory. All animal work was reviewed and approved by the University of Glasgow Animal Welfare & Ethical Review Board (AWERB), and carried out in accordance with UK Home Office regulations in line with Animals (Scientific Procedures) Act 1986, the European Directive 2010/63/EU and ARRIVE guidelines (Kilkenny et al., 2010). Mice were bred and maintained at the BICR Animal Facility. Ear notching and general maintenance (food, water and housing) was carried out by the BICR Biological Services Unit. Experimental cohorts and breeding stocks were routinely checked for health concerns. Pre-neoplastic mice were age ≤ 8 weeks old. Mice were humanely culled using rising concentration of CO_2_ at end points for the model, which include enlarged spleen (swelling in the abdomen & high gait), thymic lymphoma (difficulty breathing - rapid, shallow breathing, panting), lymph node enlargement >15 mm (commonly axillary, brachial, cervical, inguinal), reduced mobility, weight loss (pinched at the shoulders). Animals were humanely culled using Schedule 1 techniques as stipulated in the United Kingdom Home Office project licence. For routine genotyping, all animals were ear notched at weaning and samples sent to Transnetyx (Cordova, TN, US) for analysis. The *Rock1 NC* mice (Rock1^tm1.1Bicrq^) are available upon reasonable request to the corresponding author.

### Tissue collection and fixation

Animals were euthanized by carbon dioxide inhalation; weights were recorded, and blood collected for haematology or biochemistry analysis. Necropsy was performed immediately to prevent tissue autolysis. All lymphoid organs including the thymus, spleen and lymph nodes were harvested and weights recorded. Tissue were either fixed in 10% neutral buffered formalin for histological analysis or maintained in PBS until analysis. Fixed tissues were then processed, paraffin embedded and sectioned by the Histology Service at the Beatson Institute.

### Isolation of lymphocytes from tumours

Tumours (thymus, spleen or lymph nodes) were disaggregated aseptically in 4-5 mL of PBS (phosphate buffered saline; 170 mM NaCl, 3.3 mM KCl, 1.8 mM Na_2_HPO_4_, 10.6 mM H_2_PO_4_) by pushing cells through a 70 µm cell strainer into 50 mL centrifuge tubes. Lymphocytes were isolated on a Ficoll-Paque density gradient by layering disaggregated cell suspensions on 5 mL of Ficoll-Paque PLUS in 15 mL centrifuge tubes. Tubes were centrifuged at 3000 rpm for 10 minutes, and the interphase layer containing live lymphocytes was extracted and washed in 5-10 mL of RPMI 1640 (with 10% Fetal Bovine Serum (FBS), 2 mM L-glutamine, 100 U/mL Penicillin, 100 µg/mL StreptoMycin, 50 µM 2-mercaptoethanol) medium and centrifuged at 1500 rpm for 5 minutes. The supernatant was discarded, and cells were resuspended in 5-10 mL of RPMI 1640 medium. A viable cell count was carried out either with a Coulter Counter or by trypan blue exclusion with a haemocytometer. Cells were frozen at 5-10 x 10^6^ cells/vial in freezing medium (10% DMSO and 30% FBS in RPMI 1640 medium) for immunophenotyping. For tumour cell line establishment, cells from each mouse were cultured in 25 cm^2^ tissue culture flasks or 12 well plates at 2.5 x 10^6^ cells/mL in lymphocyte medium for at least 6 passages. Cell lines were frozen at a concentration of 5-10 x10^6^ cells/vial.

For transplantation of lymphoma cells, *Eµ-Myc*-expressing cells were thawed and maintained in RPMI 1640 medium. Recipient mice were sub-lethally irradiated at 5.5_Gy, and 4_h later injected with 10^6^ cells in a final volume of 100_μL through tail veins (Latif et al., 2021). Mice were maintained for two weeks on Baytril antibiotic in their drinking water pre- and post-transplantation and maintained in filter-top cages.

### Cellular protein extraction and analysis

#### Preparation of cell lysates

Lymphoma cells were plated at 2 X 10^5^ cells per well of 12 well tissue culture plates, then the next day were treated with 10 µM H1152 for 4 h, 10 µM ABT199 plus 10 ng/mL cycloheximide for 1h, or a combination of H1152 (4 h) plus ABT199 and Chx (1 h). Cell cultures were placed on ice and conditioned medium containing apoptotic bodies was collected in cold centrifuge tubes, then ice-cold PBS was added, cells were scraped and transferred to the corresponding cold centrifuge tubes. After centrifugation at 3000 rpm for 5 minutes at 4°C, supernatants were discarded, and pellets were washed in 1 mL of PBS, followed by another centrifugation at 3000 rpm for 5 min at 4°C. Supernatants were discarded and pellets were resuspended in 200-250 µL of sodium dodecyl sulfate (SDS) lysis buffer (1% (w/v) SDS, 50 mM Tris-HCl pH 7.5), which was transferred to QIAshredder tubes and centrifuged for 1 min at 14,000 rpm. Flow-through was transferred to fresh Eppendorf tubes and frozen (either rapid freezing or overnight freezing at -80°C). Lysates were thawed on ice before being centrifuged at 14,000 rpm for 15 minutes at 4 C. Supernatants were transferred to new microfuge tubes, protein concentration determined, and samples stored at -20°C.

#### Determination of protein concentration

Protein concentration of cell lysates was determined using the bicinchoninic acid (BCA) assay. Protein standards (0.08, 0.1, 0.2, 0.4, 1 and 2 mg/ml) were prepared in 1% SDS lysis buffer using a 2 mg/mL bovine serum albumin standard stock solution. 10 μL of blank (buffer alone), protein standards and samples were added to a Greiner Bio-One 96-well plate. 1:2 dilutions of protein samples were also included. 200 μL of developing solution (50:1, bicinchoninic acid: copper sulphate solution) was added to each well and incubated at 37° C for 30-60 minutes. Absorbance was measured using Molecular Devices Microplate Reader and sample concentrations determined from the standard curve.

#### SDS-polyacrylamide gel electrophoresis

SDS-polyacrylamide gel electrophoresis (PAGE) was used to separate protein samples based on molecular weight. Protein samples were first diluted in 6x loading buffer (300 mM Tris-HCl pH 6.8, 30% glycerol, 6 mM EDTA, 10% SDS, 60 mM Dithiothreitol (DTT), 0.12 mg/mL bromophenol blue) and heated at 95°C for 5 minutes. After heating, samples were briefly centrifuged and loaded on 4-12 % NuPAGE Bis-tris gels alongside Prestained Protein ladder molecular weight marker. Gels were run in tanks containing 1x NuPAGE MOPS/MES SDS running buffer at 100 V for the first 10 minutes, and then increased to 160 V and run until the tracking dye front reached the lower end of the gel. Afterwards, gels were used for western blotting.

#### Western Blotting

Following protein separation, gels were used for western blotting. Proteins were transferred to PVDF membranes using a Bio-Rad Mini Trans-Blot Cell and transfer buffer containing 20% methanol. Briefly, prior to transfer, PVDF membranes were pre-wet in methanol for 15 seconds and then washed in transfer buffer. Sponges and Whatman papers were also pre-wet and then stacked with the gel and membrane. All transfers were run at 100 V for 1 hour using an ice pack in the apparatus. Successful transfer and equal loading of proteins was confirmed by staining membranes with Ponceau staining solution. After the Ponceau stain was washed off using Tris buffered saline (TBS: 20 mM Tris-HCl pH 7.5) with 0.1% (v/v) Tween-20 (TBS-T), the membrane was blocked with 5% (w/v) milk powder for 1 hour. Primary mouse ROCK1 antibody (BD Biosciences 611136, RRID AB_398447) was diluted to the required concentration in TBS-T and membranes incubated for 2 hours at room temperature or 4° C overnight. This was followed by three washes in TBS-T for 10 minutes each. The membrane was then incubated with anti-mouse secondary antibody that had been diluted to the required concentration in TBS-T for 45 minutes at room temperature. After the blot was washed in TBS-T three times for 10 minutes each, protein bands were visualised on the LI-COR Odyssey system. Protein bands were quantified using Image Studio Version 2.1.

### Flow Cytometry

Cell suspensions were generated from mouse organs or cultured cells as follows:

#### Bone Marrow

Leg bones were dissected from the mouse and excess tissue removed using paper towel, also removing epiphysis and condyles. Bones were then placed in ice cold sterile PBS and transported to the laboratory. A hole was made in the bottom of a 0.5 mL Eppendorf tube with an 18G needle, this Eppendorf tube was placed inside a 1.5 mL Eppendorf tube. The distal end of each bone was removed with a scalpel, then the bones were placed distal end down inside the 0.5 mL Eppendorf tube and spun at 10,000 x g for 30 seconds. The bones and 0.5 mL Eppendorf tube were then discarded and the pellet within the 1.5 mL Eppendorf tube was resuspended in 1 mL red blood cell (RBC) lysis buffer (155 mM NH_4_Cl, 12 mM NaHCO_3_, 0.1 mM EDTA) for 5 minutes. Following termination or RBC lysis in 9 ml PBS, the suspension was then spun down and resuspended in PBS.

#### Blood

Blood was collected via cardiac puncture and placed into K3-EDTA tubes and mixed well to avoid clotting. Blood was then resuspended in 10 mL RBC lysis buffer and placed on a rocker for 5 minutes. Following termination of RBC lysis in 40 mL PBS, the suspension was then spun down and resuspended in PBS.

#### Thymus

Thymi were harvested, cut into small pieces, and 5 mL of thymocyte digestion mix (1 mg/mL collagenase D in RPMI) was added to the tissue in a 50 mL Falcon tube. Tissue was incubated on a shaker at 37°C for 30 minutes, then passed through 100 µM filters, followed by the addition of an equal volume RBC lysis buffer for 3-5 minutes. The RBC lysis was terminated with FBS, then cells were resuspended in PBS.

#### Immunophenotyping of lymphocytes

Frozen vials of lymphocytes isolated from *Eµ-Myc* tumours were thawed and added to 50 mL Falcon tubes containing 5 mL of RPMI 1640 (with 10% FBS), and centrifuged at 1500 rpm for 5 minutes at 4°C. Pellets were resuspended in 5 mL of FACS buffer (PBS with 5% FBS) and centrifuged at 1500 rpm for 5 minutes at 4°C. Cell pellets were resuspended in FACS buffer (volume depending on the size of the pellet). Cells were counted and divided for different staining combinations. Around 5 x 10^5^ or fewer cells were used for staining (96 well plate or Eppendorf tubes). Cells were centrifuged at 1500 rpm for 5 minutes at 4°C. Pellets were resuspended in 100 µL of FACS buffer with Fc block (1:250) and incubated for 15 min at 4°C. Meanwhile, primary antibody master-mixes were made in FACS buffer (1:500 for Biotinylated Antibodies and 1:100 for FITC/APC antibodies). After Fc block, cells were centrifuged at 1500 rpm for 5 minutes and cell pellets were resuspended in 100 µL of the relevant antibody master mix and incubated for 30 min at 4°C in the dark. Unstained and single-stained sample controls were also included. Cells were washed twice with FACS buffer. Samples containing biotinylated antibodies were resuspended in 100 µL of Streptavidin APC (1:500 in FACS buffer) and incubated for 30 min at 4°C in the dark. Cells were washed again and finally resuspended in FACS buffer containing 7AAD (1:1000) for FACS analysis.

#### Cell staining

2.5 x 10^6^ cells were stained in a volume of 100 µL unless otherwise stated. Cell suspensions were incubated in Zombie NIR live/dead dye at 1:250 in PBS at room temperature for 20 minutes. Suspensions were spun at 300 x g for 3 minutes and washed twice in PBS. Suspensions were then incubated in Fc block at 1:250 in FACS buffer (PBS with 2%(v/v) FBS, 0.5% (w/v) sodium azide, 0.2 mM EDTA) for 20 minutes at room temperature then washed twice in FACS buffer. Cells were then stained with primary antibody cocktail for 30 minutes at room temperature. If more than one antibody conjugated to a Brilliant Violet fluorophore was used in the primary antibody cocktail, the primary stain was performed in Brilliant Violet Stain Buffer, otherwise the primary stain was performed in FACS buffer. Cells were then spun down and washed twice in FACS buffer. Secondary staining containing secondary antibodies and/or streptavidin fluorophore conjugate was then performed in FACS buffer at room temperature for 30 minutes. Cells were again spun down and washed twice in FACS buffer. Cells were then resuspended in FACS buffer and an equal volume of 4% paraformaldehyde in PBS (final concentration 2%) was added for fixation.

Where intracellular staining was performed, fixation and permeabilization were undertaken with BioLegend True-Nuclear Transcription Factor Buffer kits. Cells were fixed for 1 hour in 1 mL True-Nuclear 1X Fix Concentrate in the dark. 2 mL 1X Perm Buffer was added, and samples spun down at 300 x g for 5 minutes. Samples were further washed in Perm Buffer, spun down and the supernatant discarded. Samples were resuspended in intracellular antibody cocktail in Perm buffer and incubated at room temperature for 30 minutes. Samples were spun down, supernatant discarded and resuspended in Perm buffer twice. Specific staining panels are summarised in the table below. Samples were analysed and data collected using an Attune NxT acoustic focusing flow cytometer. Post-acquisition analysis was performed using FlowJo v10 software.

**Table 1.**
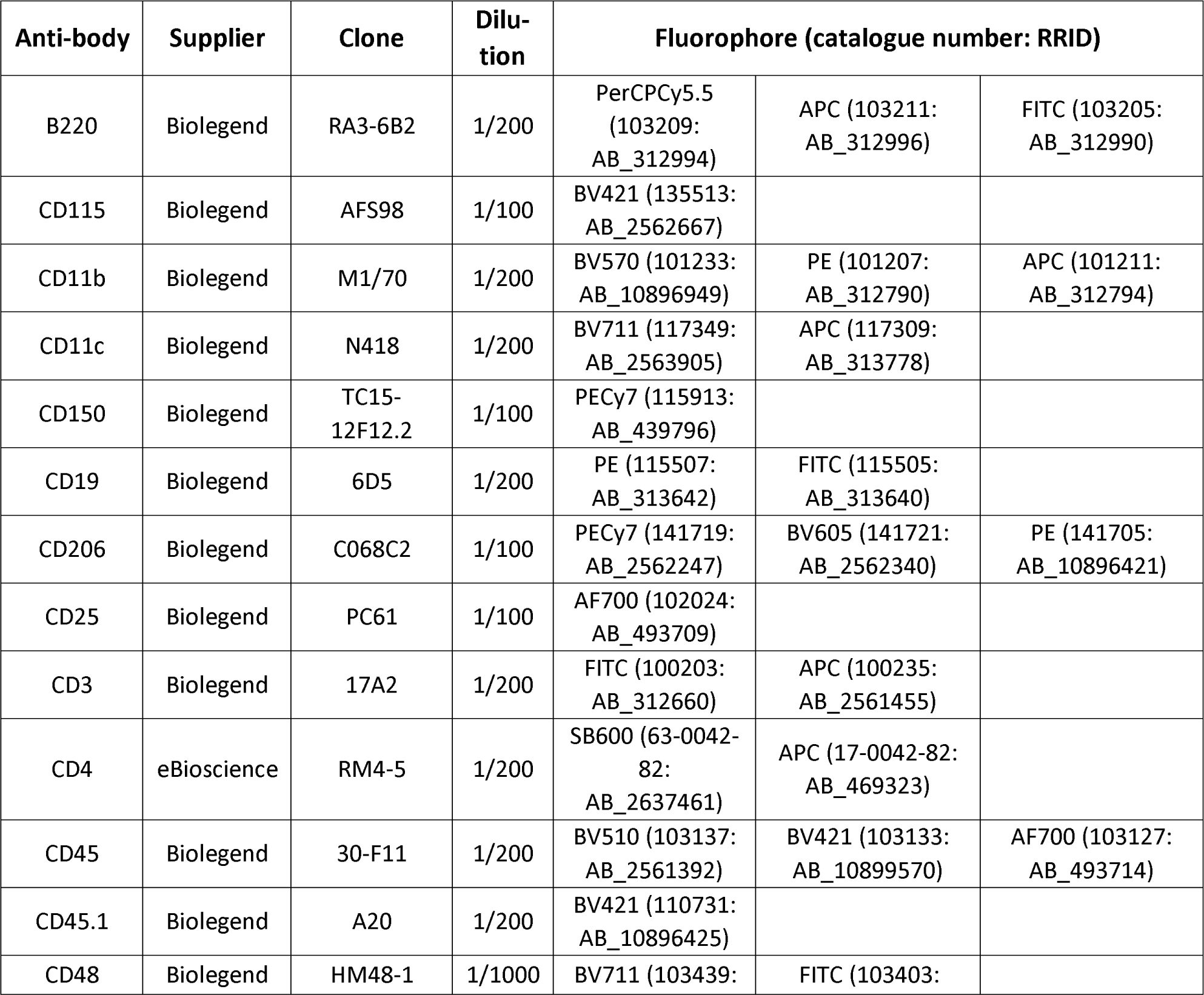
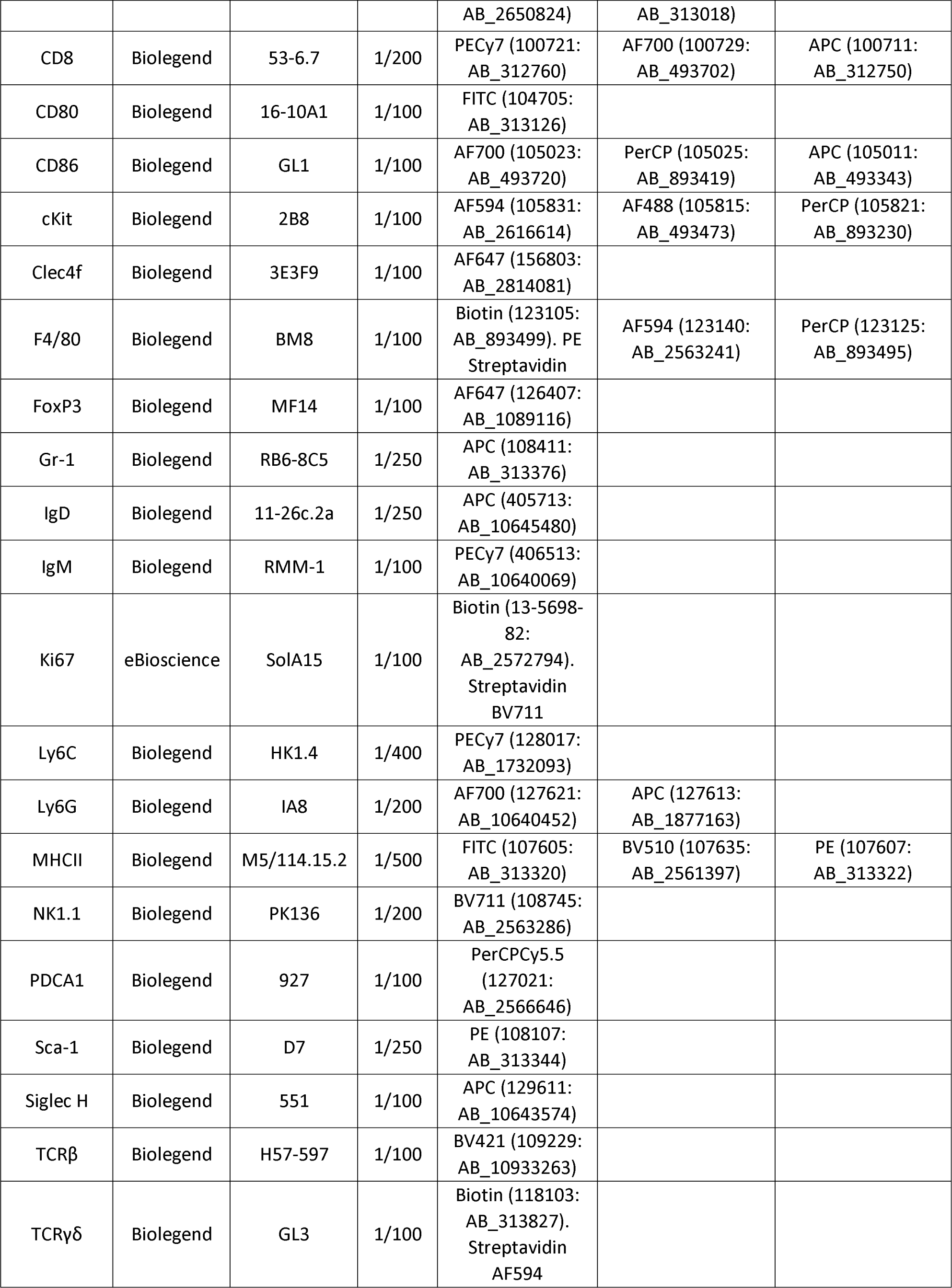
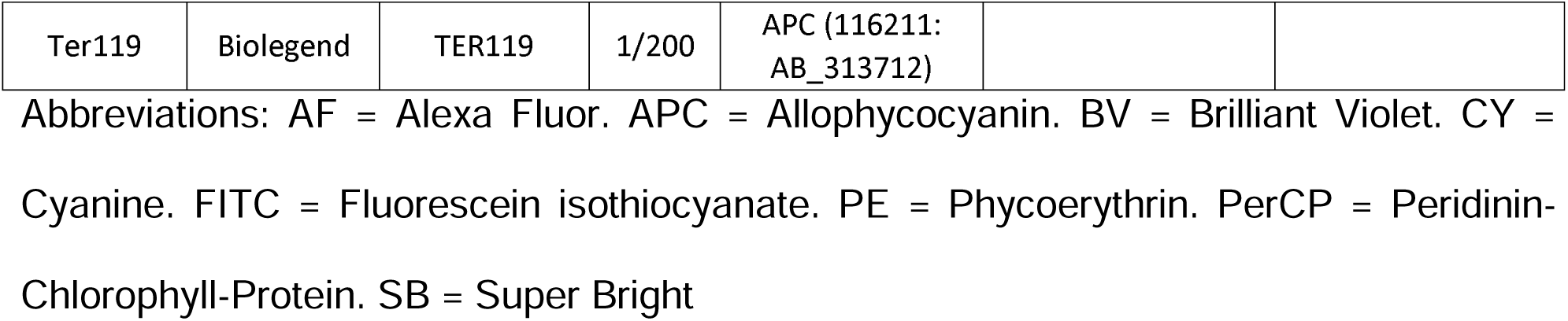
Flow cytometry antibodies.

### Clinical Pathology

#### Haematology

Mice were euthanized by carbon dioxide inhalation and blood samples were collected via cardiac puncture in potassium-EDTA tubes. Samples were immediately sent to the University of Glasgow Veterinary School Clinical Pathology Lab for complete haematology analysis. Complete blood counts were obtained using an ADVIA 120 analyser (Siemens, Frimley, UK) using 150 µL of EDTA anti-coagulated blood. Air-dried blood smears were prepared and fixed in methanol before staining with the Romanowsky stain (May-Grünwald-Giemsa). Smears were then examined by microscopy to verify analyser results, assess cell morphology and polychromasia, and a 200-cell manual white cell differential evaluation was also performed.

### Histology

Paraffin-embedded tissue sections of 4 μm thickness were cut and routinely stained with haematoxylin and eosin (H&E) by the Beatson Institute’s Histology Service. Slides were rinsed in distilled water and then stained with haematoxylin for 1 minute. After rinsing in distilled water, the slides were transferred to Scots tap water for 1 minute or until blue colour developed to the desired intensity. Slides were rinsed in distilled water and the sections were dehydrated using increasing concentrations of ethanol (1 wash for 3 minutes in 70% ethanol, 2 washes of 3 minutes each in 100% ethanol). Slides were then cleared by three washes in xylene for 3-minutes each, before mounting with DPX. Sections were analysed using bright field microscope (Olympus BX51) and representative images were obtained at different magnifications.

### Micro-CT

Dissected femurs were scanned and analyzed with a Bruker micro computed tomography Skyscan1172 at the University of Dundee according to the manufacturer’s instructions.

### Primary mouse macrophages

Mice were culled by increasing CO_2_ concentration followed by cervical dislocation. The external fur was sterilised with 70% ethanol and abdominal fur/skin removed exposing the peritoneal membrane. Following this 5 mL of sterile PBS was injected into the peritoneal cavity using a 23G needle. Each mouse was gently rocked to ensure circulation of PBS to all areas of the peritoneal cavity. Each mouse was placed over a 10 cm tissue culture dish and the peritoneal membrane was cut, draining the PBS. The cell suspension was aspirated into a sterile universal tube, and the dish washed with 10 mL sterile PBS, also placed into the universal tube. The cell suspension was then placed on ice and transferred to the laboratory. The cell suspension was filtered through a 70 µM strainer then spun at 300 x g for 5 minutes. The supernatant was discarded and the pellet resuspended in complete macrophage medium (RPMI 1640, 10% FBS (v/v), 2 mM L-glutamine, 100 U Penicillin, 0.1 mg/mL Streptomycin). Flow cytometric analysis confirmed the enrichment of macrophages (CD11b^+^, F4/80^+^) following 24 hours in culture.

For analysis of macrophage migration, primary macrophage cells were plated at 60,000 cells/well in 48 well plates in 200 µL complete medium. Cells were left overnight to attach to plates, washed twice in PBS then resuspended in complete medium. Imaging was performed on the Incucyte Zoom time lapse microscope. Post-acquisition analysis was performed using Trackmate, and then Chemotaxis and Migration software plugins in the FIJI image processing package.

For analysis of phagocytosis, primary macrophages were plated in 12 well tissue culture plates at 5 X 10^5^ cells/well. FluoSpheres (2 um diameter; Invitrogen) were opsonized with complete serum for 1-hour at 4°C. Medium was aspirated from the wells containing macrophages, then medium containing beads at 5 X 10^6^ beads/well was added, incubated for 2 hours, then phagocytosis was halted by placing cells on ice. Wells were washed once with PBS, cells were scraped, filtered through 40 µm diameter pore filters and analyzed by FACS.

### Zymosan induced peritonitis

Zymosan (SigmaAldrich) was dissolved in sterile PBS by vigorous shaking to allow even distribution of zymosan particles at a concentration of 2 mg/mL. Mice were subjected to peritoneal injection with 500 µL zymosan solution (1 mg per mouse). For peritoneal lavage, 5 mL of ice-cold lavage buffer (PBS with 2 mM EDTA) was injected into the peritoneal cavity. Mice abdomens were gently massaged to dislodge cells before collecting the fluid into ice cold collection tubes, which was then passed through 70 µm diameter pore filters. The lavage fluid was centrifuged at 1500 rpm for 8 minutes, then the cell pellet was re-suspended in FACS buffer for flow cytometric analysis.

### Statistical analysis

Statistical significance of differential findings between experimental groups was determined by performing appropriate tests using GraphPad Prism 10. The test used and sample sizes for analysis are stated in the corresponding figure legends. No animals were excluded from the analysis. Within each genotype, mice were randomly assigned to experimental groups or procedures. Experimenters were not blinded to group allocation, samples processed or evaluated by outside services (*e.g.* clinical pathology, histology) were treated blindly.

## Acknowledgements

This study was funded by Cancer Research UK institutional funding to the CRUK Beatson Institute (A10419, A17196), to the Blyth lab (A29799), and to the Olson lab (A18276). Additional funding to M.F.O from the Canadian Institutes of Health Research (PJT-169125), Natural Sciences and Engineering Research Council of Canada (RGPIN-2020-05388) and Canada Research Chairs Program (950-231665). Thanks to the Cancer Research UK Beatson Institute Biological and Histology Services, as well as the University of Glasgow Veterinary School Clinical Pathology Laboratory.

## Data availability

Data generated during this study are presented in the figures. No datasets were generated or used in this study.

